# Evolutionary and biological mechanisms underpinning chitin degradation in aquatic fungi

**DOI:** 10.1101/2024.02.10.579206

**Authors:** Nathan Chrismas, Kimberley Bird, Davis Laundon, Poppy Hesketh-Best, Chloe Lieng, Michael Cunliffe

## Abstract

Fungal biology underpins major processes in ecosystems. The Chytridiomycota (chytrids) is a group of early-diverging fungi, many of which function in ecosystems as saprotrophs processing high molecular weight biopolymers, however the mechanisms underpinning chytrid saprotrophy are poorly understood. Genome sequences from representatives across the group and the use of model chytrids offers the potential to determine new insights into their evolution. In this study, we focused on the biology underpinning chitin saprotrophy, a common ecosystem function of aquatic chytrids. The genomes of chitinophilic chytrids have expanded inventories of glycoside hydrolase genes responsible for chitin processing, complemented with bacteria-like chitin-binding modules (CBMs) that are absent in other chytrids. In the model chitinophilic saprotroph *Rhizoclosmatium globosum* JEL800, the expanded repertoire of chitinase genes is diverse and almost half were detected as proteins in the secretome when grown with chitin. Predicted models of the secreted chitinases indicate a range of active site sizes and domain configurations. We propose that increased diversity of secreted chitinases is an adaptive strategy that facilitates chitin degradation in the complex heterologous organic matrix of the arthropod exoskeleton. Free swimming *R. globosum* JEL800 zoospores are chemotactic to the chitin monomer N-acetylglucosamine and accelerate zoospore development when grown with chitin. Our study sheds light on the underpinning biology and evolutionary mechanisms that have supported the saprotrophic niche expansion of some chytrids to utilise lucrative chitin-rich particles in aquatic ecosystems and is a demonstration of the adaptive capability of this successful fungal group.

## Introduction

Fungi are leading drivers of ecosystem processes, including the degradation of high-molecular weight, often recalcitrant, biogenic organic polymers. As high-molecular weight material is too large to be directly absorbed across fungal cell walls, substrates must first be degraded in the extracellular environment. This process is carried out by enzymes secreted by the fungus into the environment before subsequent breakdown products are absorbed via osmotrophy (Richards and Talbot 2018). Our understanding of the biological mechanisms underpinning fungal saprotrophy is largely based on plant cell wall decomposition by terrestrial Ascomycota and Basidiomycota, the evolution of which is synonymous with the successful expansion of the kingdom through exploitation of diverse ecological niches (Berbee et al. 2017, Nagy et al. 2017, Naranjo-Ortiz and Gabaldón 2019).

The Chytridiomycota (chytrids) is a group of early-diverging fungi that are typically unicellular, developing into a sporangium with substrate-attaching and feeding rhizoids (Fig. 1a). Within mature sporangia, multiple uniflagellate zoospores form that lack a cell wall and are usually motile when released into the environment. Once a suitable substrate is found by swimming zoospores they attach and encyst, developing into a new sporangium with rhizoids to repeat the life cycle (Medina and Buchler 2020). Chytrids broadly occupy two major functional roles in ecosystems, either as (i) parasites, mainly of algae with the exceptions of a few amphibian pathogens, or (ii) saprotrophs that degrade a range of recalcitrant organic biopolymers. A recent phylogenomic study has proposed that saprotrophy is a derived trait in chytrids and that the ancestral trait is algal parasitism (Thomé et al. 2023). The biological mechanisms underpinning aquatic chytrid saprotrophy are currently poorly understood including in an evolutionary context (Laundon and Cunliffe 2021).

**Figure 1.**
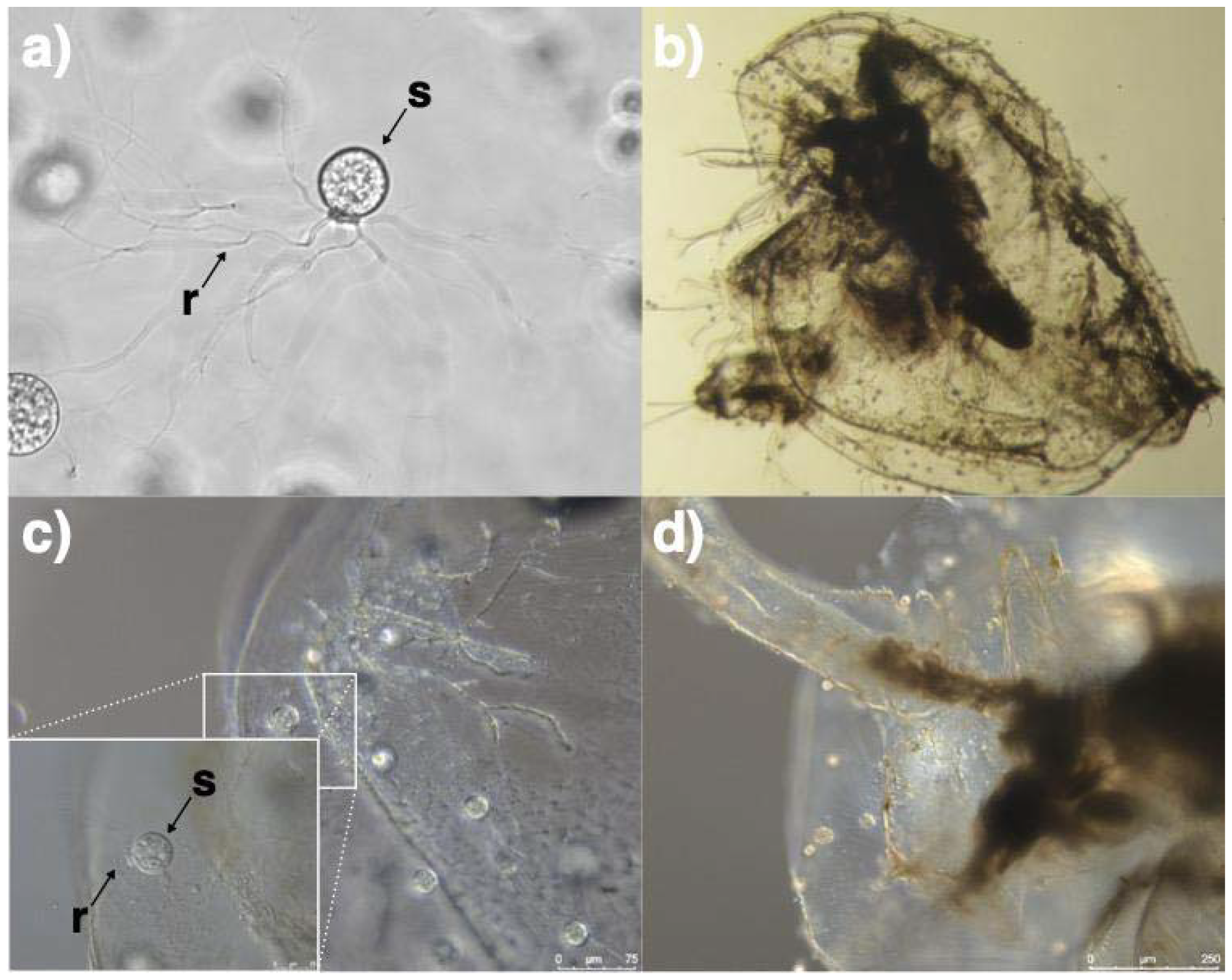
a) A *Rhizoclosmatium globosum* JEL800 cell with branching rhizoids (r) used for attaching to substrates and sporangium (s) inside which develop multiple zoospores. b-d) *Rhizoclosmatium globosum* JEL800 attached to the *Daphnia* exoskeleton across various levels of magnification. Note rhizoids (r) penetrating the exoskeleton surface.

Chitin-based particles such as arthropod exoskeleton remains (exuviae) are prominent aquatic chytrid habitats (Sparrow 1960) (Fig. 1b-d). Chitin is a polymer of N-acetylglucosamine (GlcNAc) and is the second most abundant polysaccharide on Earth after cellulose. Chitin degradation is carried out by chitinases that either remodel cellular chitin (e.g. in the fungal cell wall), or externally process chitin substrate (Duo-Chuan 2006). Fungal chitinases belong to the glycoside hydrolase family 18 (GH18) of carbohydrate-active enzymes (CAZymes) (Funkhouser and Aronson 2007) and act by introducing random breaks into crystal chitin (endochitinases) or cleaving terminal ends of GlcNAc polymers (exochitinases). GH18 chitinases are classified into two main related groups (Goughenour et al. 2021). Group AC are typically exochitinases with a deep, tunnel shaped substrate-interacting groove whereas Group B are typically endochitinases with a shallow, open groove. GH18 domain-containing proteins can also include carbohydrate binding modules (CBMs) belonging to ChtBD3/ChiC_BD domain families, which increase substrate binding affinity. Following primary chitinolytic activity by GH18s, chitin-derived GlcNAc is further processed by GH20 family β-*N*-acetyl-D-hexosaminidases.

The aims of this study were to determine the biological mechanisms underpinning chytrid chitin saprotrophy by focusing on the following specific questions: (i) what is the underlying molecular machinery (i.e. genes, mRNA and proteins/enzymes) involved in chitin saprotrophy, and (ii) what chitin-associated behaviour is shown by swimming zoospores. Our approach was to compare the reference genome sequences from chitinophilic chytrids with representatives across the Chytridiomycota that occupy a range of different ecological niches, combined with the use of the model saprotroph *Rhizoclosmatium globosum* JEL800 to determine new insights into chytrid biology. *Rhizoclosmatium globosum* is an aquatic chytrid found prevalent in nature attached to chitin-rich arthropod exuviae (Sparrow 1960). The strain *R. globosum* JEL800 is an emerging model for biological and evolutionary studies of early diverging fungi (Laundon et al. 2020, Venard et al. 2020, Prostak et al. 2021, Galindo et al. 2022, Laundon et al. 2022) that was isolated by chitin baiting (Joyce Longcore, pers. comm.) and grows readily on chitin as a sole carbon source (Laundon et al. 2020).

## Results

### Expansion of genes encoding chitin degradation enzymes in chitinophilic chytrids

The abundance of GH18 chitinase encoding genes in the reference genomes of 36 chytrids across the Chytridiomycota was compared, including the Chytridiomycetes Chytridiales (n=7), Caulochtriales (n=1), Cladochytriales (n=2), Polychytriales (n=1), Rhizophydiales (n=5), Spizellomycetales (n=6), *Blyttiomyces helicus incertae sedis* (n=1), as well as Monoblepharidomycetes (n=2) and Neocallimastigomycetes (n=11) that occupy a diverse range of ecological niches (Fig 2ai, Supplementary File, Supplementary Figure 1).

**Figure 2.**
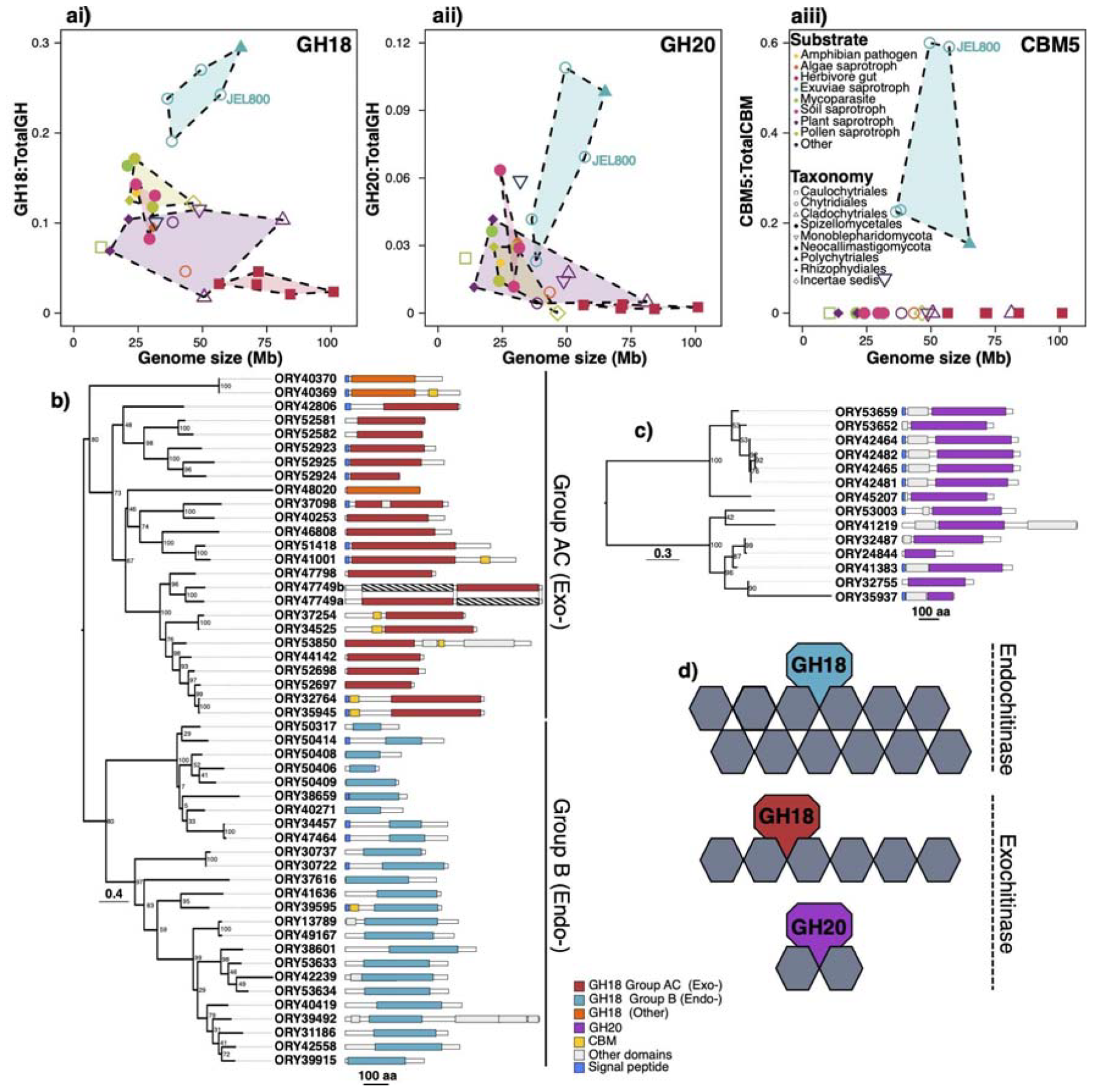
ai) Comparison of ratios of glycoside hydrolases GH18 chitinases and aii) GH20 β-*N*-acetyl-D-hexosaminidases with total GH encoding genes, and aiii) carbohydrate binding module CBM5 genes with total CBM encoding genes and genome sizes of 36 members of the Chytridiomycota that occupy diverse ecological niches and utilise a range of substrates (full labels in Supplementary Figure 1). Point colours indicate the substrate strains were isolated from, convex hulls link strains isolated from the same substrate; point shapes indicate order. Maximum likelihood phylogenies of *Rhizoclosmatium globosum* JEL800 b) GH18 and c) GH20 domains showing architecture of GH domain-containing genes. Genes encode for proteins with either d) endo-(GH18) or exochitinase (GH18, GH20) activity.

All chytrids associated with arthropod exuviae in nature and isolated by chitin-baiting belong to the Chytridiales except *Polychytrium aggregatum* JEL109 from the sister order Polychytriales. All the chitinophilic chytrids i.e. *R. globosum* JEL800, *Obelidium mucronatum* JEL802, *Chytriomyces hyalinus* JEL632, *Chytriomyces* sp. MP 7 and *P. aggregatum* JEL109 showed the highest enrichment of GH18 encoding genes (Fig 2ai, Supplementary Fig 1). All other chytrids, i.e. occupying non-chitin associated niches, had relatively lower GH18 gene abundances with the plant-processing Neocallimastigomycetes found in herbivore guts and plant-baited *Cladochytrium replicatum* JEL0714 lowest (Fig 2ai, Supplementary Fig 1). A similar pattern of increased gene frequency was observed for GH20 encoding genes, particularly for the chitinophilic *R. globosum* JEL800, *O. mucronatum* JEL802 and *P. aggregatum* JEL109 (Fig. 2aii, Supplementary Fig 1).

### Bacteria-like chitin binding module genes in chitinophilic chytrids

The abundance of genes containing carbohydrate binding module 5 (CBM5) domains in the genomes of the chitinophilic chytrids was distinctive, with *R. globosum* JEL800 and *O. mucronatum* JEL802 particularly high (Fig. 2aiii, Supplementary Fig 1). Although GH18 and GH20 encoding genes were present but at lower abundances in the genomes of non-chitin associated chytrids (Fig. 2 ai-aii), genes containing CBM5s were only detected in the genomes of chitinophilic chytrids except for the Monoblepharidomycete *Hyaloraphidium curvatum* SAG235-1. In most cases, BLAST searches of the CBM5 domains from chitinophilic chytrids revealed similarity with equivalent bacterial ChtBD3/ChiC_BD domains (Supplementary File), many of which were found in bacteria with known chitin-degrading capacity (e.g. *Chitinilyticum aquatile, Paenibacillus chitinolyticus, Chitinolyticbacter albus*). Only three genes, one in *Chytriomyces* sp. MP71 (KAI8611252) and two in *Polychytrium aggregatum* JEL109 (XP_052967452 and XP_052967585), contained domains more similar to those found in non-chytrid fungi with matches to Agaricomycetes (Basidiomycota). In addition, our CBM5 domain searches matched to a further three Chytridiales genomes not included with the reference Mycocosm genomes (*Podochytrium* sp. JEL0797, *Siphonaria sp*. JEL0065, and *Physocladia obscura* JEL0513), all of which were isolated from mayfly (Ephemeroptera) exuviae (Joyce Longcore, pers. comm.).

### Rhizoclosmatium globosum GH18 and GH20 gene diversity and domain architecture

Forty-nine *R. globosum* JEL800 genes contained GH18 domains. Maximum-Likelihood phylogenetic analysis of the GH18 domains revealed two major clades that corresponded with either Group AC (predominantly exochitinases) or Group B (predominantly endochitinases) (Fig. 2b). Group AC contained domains for 24 proteins, with 21 forming a large sub-clade, with two domains appearing in ORY47749. Although ORY48020 fell within this clade, it was separated by a long branch with relatively low bootstrap support. ORY40370 and ORY40369 fell outside the main clade. Group B contained domains from 25 proteins. One clade contained domains from 16 proteins and a second clade contained a variety of GH18 domains from nine proteins with low bootstrap support for several branches within this clade.

Chitin binding domains, all of which were CBM5-type, were found on eight genes (Fig. 2b). Seven were associated with Group AC (ORY40369, ORY41001, ORY53850, ORY32764, ORY35945, ORY37254 and ORY34525) and one with Group B (ORY39595). Signal peptides associated with protein secretion were found in 17 proteins. Eleven of these were found in Group A proteins (45.8% of Group A) and six with Group B (24% of Group B). Fourteen *R. globosum* JEL800 genes were found to contain GH20 domains (Fig. 2c), Maximum-Likelihood phylogenetic analysis of which revealed two well supported clades, each containing seven genes, nine of which had associated signal peptides.

### GH18 and GH20 in the Rhizoclosmatium globosum secretome and transcriptome

A total of 23 proteins from GH18 domain-containing genes were detected in the *R. globosum* JEL800 secretome when grown on chitin as the sole source of carbon: 10 from Group AC (exochitinases), 12 from Group B (endochitinases) and one other (Fig. 3a), accounting for almost half (47%) of the GH18 genes detected in the genome. Proteins for six of the GH20 domain-containing proteins were also detected in the secretome (Fig. 3a). All GH18 and GH20 proteins detected in the secretome matched reciprocal mRNA sequences detected in the cellular transcriptome, with a positive correlation of secretome GH18 and GH20 protein abundance with mRNA fpkm (Fig. 3b). Predicted structures of the proteins encoded from GH18 domain-containing genes that were detected as secretome proteins suggests that *R. globosum* JEL800 chitinases are variable both in terms of total size and substrate binding site size (Fig. 3c and Supplementary Figures 2 and 3). Modelling of the abundant GH18 protein ORY35945 predicted a C-terminal GH18 domain and N-terminal CBM5 domain joined by a linker domain (Fig. 3d).

**Figure 3.**
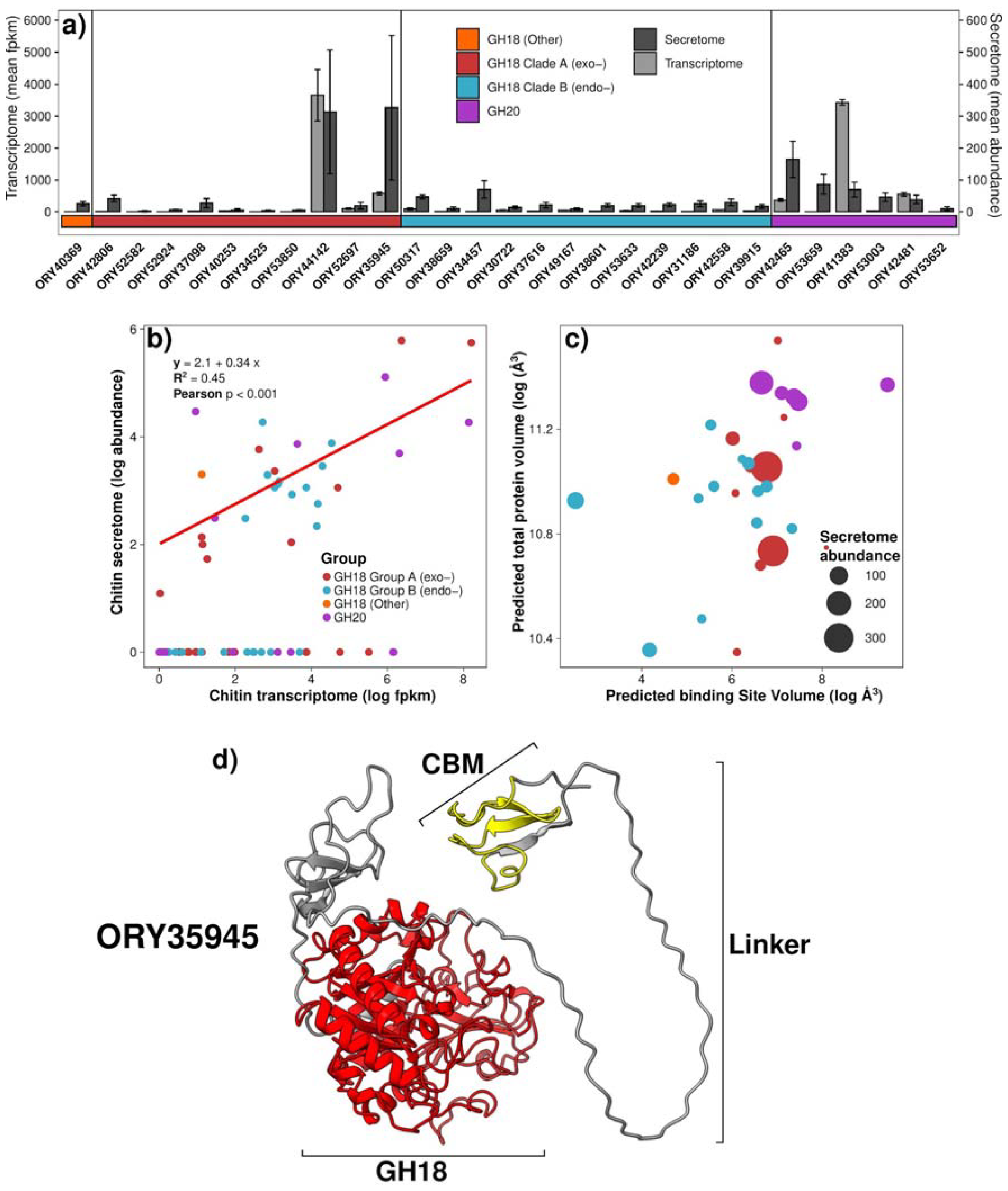
a) Secretome abundance and mRNA (fpkm) for all *Rhizoclosmatium globosum* JEL800 glycoside hydrolases present in the secretome. b) Relationship of abundance of proteins in the secretome to equivalent mRNA in the transcriptome c) Relationship of substrate binding site volume to total protein volume d) AlphaFold predicted model of ORY35945. Colours are based on conserved domains: the carbohydrate binding module domain (CBM) (amino acid residues 22 - 36) is shown in yellow and the GH18 domain (amino acid residues 203 - 596) is shown in red. Both are joined by a linker region. Signal peptide region not shown.

### Rhizoclosmatium globosum zoospore behaviour indicates adaptations to chitinophilic lifestyle

Swimming *R. globosum* JEL800 zoospores are chemotactic towards the chitin monomer GlcNAc. Targeted swimming occurred with zoospores from cultures grown with GlcNAc as the sole carbon source (Fig. 4ai) and when grown without GlcNAc in the complex medium PmTG (Fig. 4aii). Even though *R. globosum* grows with glucose as an alternative sole carbon source (Supplementary Figure 4), chemotaxis did not occur towards glucose with either GlcNAc or PmTG grown cultures (Fig. 4ai, Fig. 4aii).

**Figure 4.**
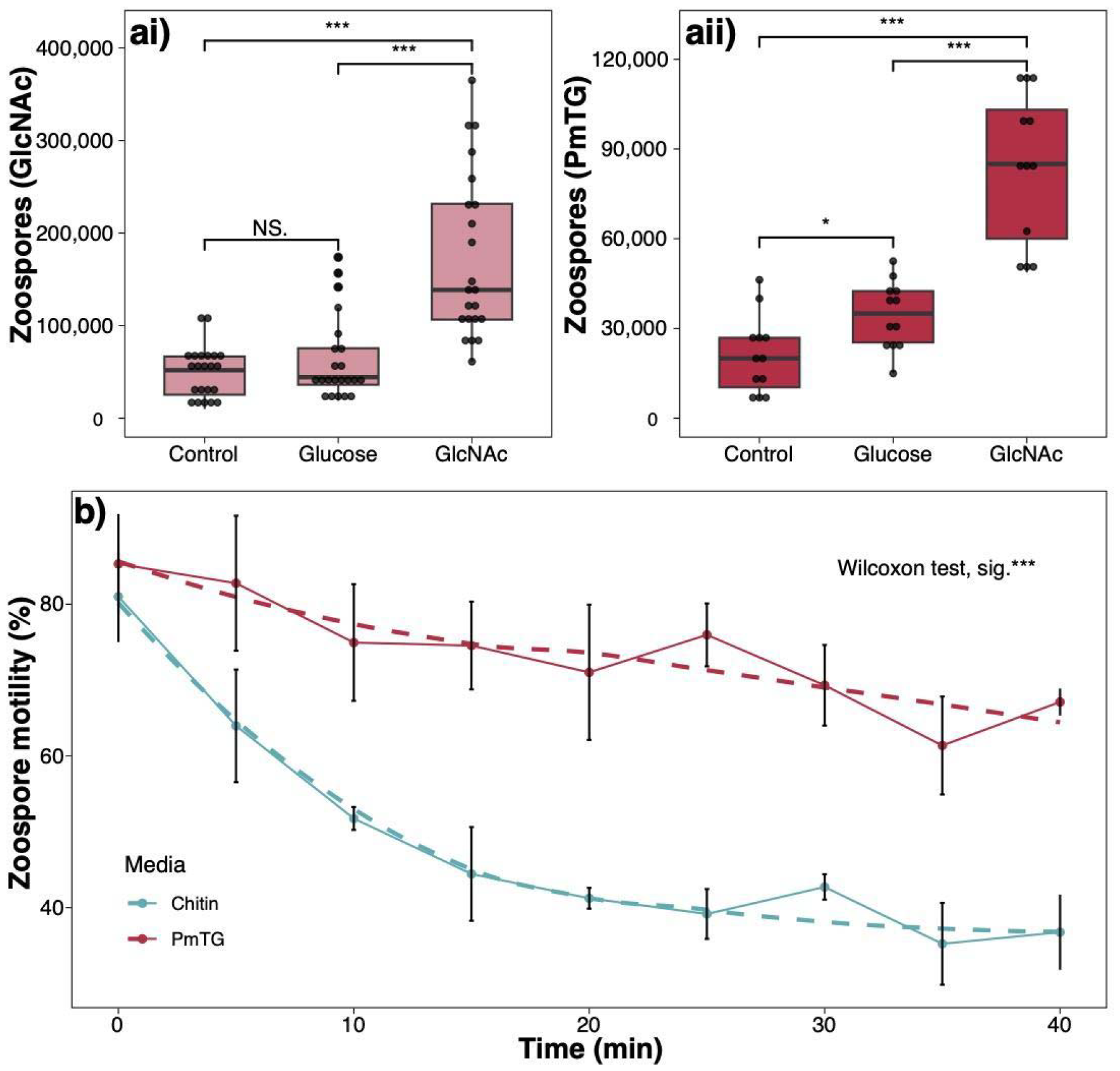
ai-ii) Chemotactic response of *Rhizoclosmatium globosum* JEL800 zoospores to the chitin monomer N-acetylglucosamine (GlcNAc) compared to glucose and no-attractant control from cultures grown with GlcNAc as the sole carbon source and in complex GlcNAc-free medium PmTG. Significance values (pairwise Wilcoxon test) are indicated as *** = p < 0.001, * = p < 0.05, NS. = not significant. Points are binned at 1/30 range of the data. b) Loss of motility in *R. globosum* JEL800 zoospores from cultures grown with chitin as the sole carbon source compared to the complex medium PmTG.

We made several attempts to determine chemotaxis in zoospores from cultures grown with chitin but were unable to conduct the assay due to apparent reduction in zoospore motility. To explore this further, we compared general motility (i.e. swimming versus stationary) of zoospores harvested from cultures grown with chitin to the motility of zoospores from cultures grown with PmTG, showing that chitin-grown zoospores rapidly lose motility after release compared to zoospores grown with PmTG which remained more motile for longer (Fig. 4b).

## Discussion

### Gene expansion and gene transfer underpin chitinophilic chytrid evolution

Here we provide genome-based evidence for two complementary mechanisms through which chitin saprotrophy has evolved in chytrids. The abundance of genes encoding GH18 chitinases and GH20 β-N-acetyl-d-hexosaminidases responsible for the breakdown of chitin and subsequent processing of chitin-derived GlcNAc respectively were increased in chitinophilic chytrids relative to non-chitinolytic chytrids. Alongside GH18 gene expansion, bacteria-like chitin-binding CBM5 domains were in the genomes of the chitinophilic chytrids, indicating that horizontal gene transfer (HGT) has taken place, potentially from chitinophilic bacteria to chitinophilic chytrids. The acquisition of chitin-binding capability likely improved the functional efficiency of chytrid chitinases to degrade extracellular chitin-based substrates and therefore has clear fitness advantages.

Lineage specific gene family expansion is a mechanism of adaptive evolution in eukaryotes and including in fungi (Lespinet et al. 2002). Expansion of hydrolase gene families related to host/substrate specificity has been reported in other fungi. In non-chytrid fungi, necrotrophic Ascomycota and Basidiomycota exhibit expansion of GH28 family pectinases (Sprockett et al. 2011), and white rot fungi including *Armillaria* spp. (Basidiomycota) have an increased complement of genes encoding cellulose- and xylan-degrading enzymes (Sipos et al. 2017). Increased copy numbers of proteases have been associated with the evolution of amphibian pathogenicity in *Batrachochytrium dendrobatidis* by facilitating attachment and penetration of host cells (Joneson et al. 2011, Farrer et al. 2017). Specifically in terms of GH18 gene expansion, as we show here in early-diverging chitinophilic chytrids, similar has been observed across other fungi including various Ascomycota (Karlsson and Stenlid 2008) and ectomycorrhizal fungi (Maillard et al. 2023).

A key feature of gene family expansion in fungi is that it facilitates exploitation of specific niches by allowing increased rates of enzyme production and functional novelty (Gladieux et al. 2014). The increased GH18 and GH20 inventory in chitinophilic chytrids may confer the ability to react quickly to substrate availability by transcribing multiple genes. Although cellular mRNA was detected for almost all of the GH18 and GH20 genes encoded within the *R. globosum* JEL800 genome, not all transcribed genes were detected in the secretome. As well as accounting for cell wall processing cellular chitinases (which are yet to be characterised in chytrids), some of this untranslated mRNA may play a role maintaining a bank of mRNA for a variety of chitinases, *R. globosom* JEL800 may be able to remain ‘primed’ for rapid protein synthesis.

Increased chitinase gene copy number in chitinophilic chytrids contrasts with chytrids occupying other niches which have reduced abundances of chitinase encoding genes. It is important to reiterate that GH18 chitinases are likely used for remodelling chitin in the chytrid cell wall based on analogous mechanisms in other fungi (Gow et al. 2017) and therefore necessary in all chytrids. Our study suggests that chitinase gene family expansion is varied within lineages, with an apparent increase in chitinase gene copy number associated with the base of the Chytridiales encompassing the closely related chitinophilic *Polychytrium aggregatum* (Polychytriales) followed by subsequent gene reduction in the plant saprotroph *Entophlyctis luteolus* (Chytridiales).

The presence of chitin-binding CBM5 domains is a distinguishing feature of chitinophilic chytrid genomes. In *R. globosum* JEL800, chitin-binding domains are present on multiple chitinases and are potentially derived from xenologs originating from chitinophilic bacteria. Horizontal gene transfer is well established in prokaryotes but has only relatively recently been determined in fungi including early-diverging fungi. Comparative assessment of herbivore gut anaerobic Neocallimastigomycetes has shown the HGT played a role in their evolution (Murphy et al. 2019). As we propose here for aquatic chitinophilic chytrids, anaerobic gut Neocallimastigomycetes HGT occurred from functionally similar bacteria that likely co-occur in the same niche i.e. anaerobes in herbivore guts that degrade plant-derived polysaccharides and is especially associated with CAZymes and extracellular polysaccharide degradation. Murphy *et al*. (2019) proposed that HGT, including the acquired ability to degrade plant-derived polysaccharides, was a key feature of the expansion of the Neocallimastigomycetes, as we propose here with the evolution of the Chytridiales. Wider comparison across fungal divisions including Mucoromycota indicates that the early diverging groups of aquatic fungi are more likely to undergo HGT compared to terrestrial taxa (Ciach et al. 2024). Further work is needed using more sophisticated bioinformatic approaches to explore HGT in chitinophilic chytrids and other saprotrophic guilds.

### Diversity in chitinase gene architecture and modelled secretome chitinase structure

Relatively few crystal structures of fungal chitinases have been characterised (Jiang et al. 2021) and prediction of specific chitinase functions is therefore difficult (Goughenour et al. 2021) but variation in binding site volume relative to overall protein volume indicates structural diversity which may be related to variation in substrate, with different enzyme activities able to account for substrate complexity (discussed further below). The high levels of secretion of Group AC exochitinases compared to Group B endochitinases suggests that cleavage of terminal GlcNAc dimers from chitin is a particularly important characteristic of the chitinophilic lifestyle.

Alongside size complexity, we also show that some chitinases are multimodular with additional domains that potentially expand the chytrids chitin degradation capability. Considering the major secreted chitinase ORY35945 (Fig. 3d), this has broadly analogous domain configurations with the cellulose degrading endoglucanase (EG11) in *Rhizopus stolonifer* (Mucoromycota) with a C-terminal catalytic domain linked with N-terminal CBM (Tang et al. 2015). The function of linkers between catalytic and binding domains has not been considered with chitinases but has been studied with cellulose and lignocellulose degrading enzymes and suggest that linkers may have different functions including general improvements in catalytic efficiency (Payne et al. 2013, Tang et al. 2015, Courtade et al. 2018), promotion of localised and repeated substrate processing (Courtade et al. 2018) and also have an additional substrate binding role (Payne et al. 2013). In terms of the chitinophilic chytrids hosting bacteria-like CBM5 domains, homologous CBM5 from the chitinophilic bacterium *Serratia marcescens* exhibits affinity specifically for α-chitin and promotes chitin degradation in the presence of other polysaccharides (Liu et al. 2023).

In nature, chitinophilic chytrids are observed growing on arthropod exoskeleton remains (Sparrow 1960) in which chitin is not available as a distinct molecule but instead packaged within a complex heterologous organic matrix. Even though exoskeleton matrices likely vary between source taxa, the general structure is based around crystalline α-chitin fibres in chitin-protein planes that are stacked in a twisted ‘plywood’ structure (Raabe et al. 2006). Taken together, we propose that the configuration diversity from varied chitinase sizes, associated bacteria-like CBM5 domains with likely α-chitin affinity and further capability caused the presence (or absence) of linkers between catalytic sites and binding domains has adaptive advantages for saprotrophy within the naturally heterologous organic matrix in which the primarily α-chitin substrate is intertwined.

### Zoospore behaviours as adaptive traits for the chitinophilic lifestyle

*Rhizoclosmatium globosum* JEL800 zoospores are chemotactic to the chitin monomer GlcNAc independent of the growth conditions of the previous generation, a trait that has obvious advantages for targeting chitin-rich substrates such as arthropod exuviae in complex aquatic environments. Chemotaxis has been shown in non-saprotrophic chytrid zoospores, particularly species that are parasites. *Batrachochytrium dendrobatidis* zoospores are chemotactic towards amphibian-produced compounds (Moss et al. 2008) and algal parasites target cell extracts of stressed hosts (Scholz et al. 2017). The biological mechanisms underpinning chemotaxis in aquatic chytrids are poorly understood (Laundon and Cunliffe 2021). It is possible that response to environmental signals could be linked to the rumposome, an enigmatic structure connecting the zoospore cell surface to the flagellar machinery that is present in *R. globosum* JEL800 zoospores (Laundon et al. 2022) and is likely a site of calcium-mediated signalling (Powell 1983). Phototaxis has also been shown in chytrid zoospores, including with *R. globosum* JEL800, facilitated by a CyclOp-fusion protein based sensory system (Galindo et al. 2022). Together, this suggests that via chemotaxis and phototaxis, swimming chytrid zoospores utilise multi-signal mechanisms to navigate the aquatic environment.

The chemotactic response to GlcNAc shown here suggests that, in nature, chitinophilic chytrid zoospores respond to signals of ‘on-going’ chitin saprotophy that is releasing GlcNAc via degradation by other organisms. Chitin degradation in aquatic ecosystems is also performed by chitinophilic bacteria that share the same habitat and potential niche as chitinophilic chytrids such as *R. globosum* (Roberts et al. 2020). It is likely that a range of chytrid-bacteria interactions are associated with chitin-rich particles, including particle colonisation, that involve the release of chemotactic signals from bacteria to chytrids and *vice versa* depending on which are the primary or secondary colonisers (Roberts et al. 2020).

When grown with chitin, we observed faster loss of next generation zoospore motility compared to non-chitin grown *R. globosum* JEL800 which remained motile for longer. Accelerated transition from a non-feeding swimming zoospore to the substrate attaching stage of the life cycle (i.e. encystment) is a possible trait that could facilitate the rapid colonisation and exploitation of ephemeral chitin-rich substrates such as mayfly exuviae in a dynamic resource-complex and competitive environment. Chytrid zoospore biology is poorly understood, including controls on the cell cycle (Medina and Buchler 2020, Laundon et al. 2022), however based on frog mucus exposure experiments with *B. dendrobatidis*, likely involve critical host/substrate chemical cues that initiate encystment via calcium signalling and actin cytoskeleton remodelling (Robinson et al. 2022). Chytrid zoospores are metabolically active, yet are transcriptionally and translationally inactive, with ribosomes held dormant (Medina and Buchler 2020). As with other chytrids, in *R. globosum* JEL800 zoospores, ribosomes are maintained dormant in a distinct cluster that only dissipates once the cell transitions to the germling stage when transcription and translation are initiated (Laundon et al. 2022). Therefore, we propose the molecular mechanisms underpinning the zoospore behavioural traits that we report here that likely support chitin utilisation are maternally derived during zoosporogenesis and combine independent (i.e. GlcNAc chemotaxis) and dependent (i.e. accelerated zoospore-germling transition) of maternal growth conditions (i.e. with or without chitin).

## Conclusion

Our study has shown mechanisms that underpin and have likely supported the evolution of chitin-based saprotrophy in chytrids. We propose that the complementary expansion of existing GH18 and GH20 genes, probably from cell wall remodelling functions, alongside niche-specific novel gene transfer of GH18-associated chitin-binding CBM5 domains, potentially from functionally similar bacteria, facilitated chitin saptrophy evolution. Expansion of GH18 genes, acquisition of CBM5 domains and linkers, allowed the diversification of secreted chitinases that may facilitate utilisation of chitin within the complex arthropod exoskeleton matrix and, along with substrate-dependent zoospore behavioural traits, determined the widespread and successful exploitation of the chitin-based particle niche in aquatic ecosystems.

## Methods

### Comparison of GH18, GH20 and CBM5 encoding genes across chytrid genomes

Counts of predicted GH18, GH20 and CBMs genes were retrieved from Chytridiomycota reference genomes from Mycocosm (https://mycocosm.jgi.doe.gov/, accessed 10/2023) (Supplementary File, Supplementary Figure 5). Conserved domains in *R. globosum* JEL800 GH18 and GH20 genes were identified using CD-search (Yang et al. 2020) and aligned separately in MAFFT (Katoh and Standley 2013) using E-INS-i before maximum likelihood phylogenetic analysis was carried out using the CIPRES (Miller et al. 2010) implementation of RaxML 8.2.10 (Stamatakis 2014) using the CAT model and the BLOSUM62 matrix for divergent protein sequences with automatic bootstrapping. Domain trees were visualised using FigTree v.1.4.4 (http://tree.bio.ed.ac.uk/software/figtree/). GH18 domains were further subclassified by comparison with a phylum-wide sampling of fungal GH18 domains (Junges et al. 2014) (Supplementary Figure 5). Presence of signal peptides in each gene were predicted using the SignalP 4.1 webserver (Petersen et al. 2011). Complete protein domains were plotted using genoplotR (Guy et al. 2010). Chitin binding domains (ChtBD3/ChiC_BD) in genes annotated as CBM5 in Mycocosm for all CBM5-containing lineages were identified using CD-search. Where domain homology was identified, domains were searched against GenBank using BLASTp (e-value cutoff = 0.00001) excluding hits for the chitinophilic chytrids in this study.

### Rhizoclosmatium globosum JEL800 maintenance, growth and microscopy

*Rhizoclosmatium globosum* J EL800 was routinely maintained on peptidised milk, tryptone, and glucose (PmTG) agar plates (Barr 1986) at 22°C in the dark. For experimental analysis, *R. globosum* JEL800 was also grown with modified Bold’s Basal Medium (BBM) (Laundon et al. 2020) diluted 10-fold and with the addition of 0.25 g ammonium sulfate and 0.5 mL f/2 vitamin stock in 1 L (Guillard 1975). To produce samples for transcriptome and secretome analysis, *R. globosum* was grown in 50 mL BBM with 10mM chitin in 250 mL Erlenmeyer flasks inoculated with ∼14,000 zoospores and shaking at 70 rpm. Cultures were harvested after 7 days growth by transferring contents to a 50 mL centrifuge tube and centrifuging at 4,000 G for 10 min. Cell pellets were snap frozen with liquid nitrogen. The supernatant was removed and Halt™ Protease & Phosphate Inhibitor Cocktail (Thermo) added at 5 µL / 50 mL with mixing by inversion. Protease treated supernatant (∼ 50 mL) was concentrated to ∼ 250 µL in 15 mL batches at 4,000 G for 20 min each (total ∼ 1 hour) using 10 kDa Amicon® Ultra 15 mL Centrifugal Filters (Millipore). Concentrated supernatant was transferred to a 3 kDa Pierce™ Protein Concentrator PES 0.5 mL (Thermo) and centrifuged for 20 min at 15,000 G to a final volume of ∼ 100 µL. All centrifuge steps were carried out at 4 °C and samples stored at -80 °C.

To screen a range of alternative non-chitin carbon sources used by *R. globosum* JEL800 for growth, mature cultures on PmTG agar plates were sporulated using 4 mL dH_2_O for ∼ 2 hours at room temperature. Zoospores were then harvested and passed through a 10 µm sieved before being washed 3 times in carbon free BBM by centrifuging at 4000 G for 10 min at 4 °C and re-suspended in carbon free BBM. Biolog PM 1 and PM 2A Carbon Utilisation Assay plates were inoculated with 150 µL per well at a final density of 1 x 10^3^ zoospores per well. Assay plates were incubated at 22 °C and absorbance at 405 nm (OD_405_) was measured periodically using a plate reader (CLARIOstar).

To demonstrate *R. globosum* JEL800 growth on an arthropod exoskeleton, *Daphnia* were acquired from a local aquarium supplier and cleaned by incubation in 10% sodium dodecyl sulphate solution for 30 mins, followed by 100% ethanol for 1 hour and then autoclaved in ddH_2_O. Carapaces were washed twice in sterile ddH_2_O between each step. Culture dishes containing 3mL BBM and one carapace were inoculated with 500 µl zoospore suspension (1.03 x 10^6^ zoospores.ml^-1^) and incubated at 22°C. Chytrids growing on carapaces were imaged with DIC using a Leica Dmi8 microscope (Leica, Germany).

### Transcriptome and secretome analysis

RNA was extracted from the cell pellets using the RNeasy Micro kit (Qiagen) according to the manufacturer’s instructions. Pellets were thawed in 2 mL RLT lysis buffer (Qiagen) containing 10 μL mL^-1^ of 2-mercaptoethanol, incubated at room temperature for 5 minutes and mixed periodically using a vortex. Cell debris was removed by centrifuging at 4000 G for 1 min, the lysate was recovered through a QIA shredder (Qiagen). An equal volume of 100% ethanol was added to the homogenised lysate before being transferred in batches to the RNeasy extraction column including on column DNase digestion step using the RNase-Free DNase set (Qiagen). RNA was quantified using a NanoDrop 1000 spectrophotometer (Thermo) and the RNA BR assay kit (Invitrogen) on the Qubit 4 fluorometer (Invitrogen).

RNA quality was assessed using the RNA 6000 Nano kit total RNA assay (Agilent) run on the 2100 Bioanalyzer instrument (Agilent). Sequencing was carried out using Illumina NovaSeq 6000 technology and assembled with the Novogene WTS pipeline. Raw reads are deposited in SRA under BioProject PRNJA1076503. GH18 and GH20 domain containing genes were identified in the assembled transcriptome by performing blastn searches for each gene identified in the *R. globosum* JEL800 genome (e-value cutoff = 0.00001). Concentrated supernatant samples (i.e. secretome) were sent to University of Bristol Proteomics Facility for comparative proteomics using tandem mass tagging (TMT). Identified peptides were cross referenced with predicted proteins from mRNA identified within the transcriptome.

### Protein modelling from gene sequences

GH18 and GH20 models were retrieved from the AlphaFold Protein Structure Database (Jumper et al. 2021, Varadi et al. 2022). Total model volumes were predicted using VADAR 1.8 (Willard et al. 2003) and binding site volumes were predicted using CASTp 3.0 (Tian et al. 2018) with default settings. The complete model for ORY35945 including the GH18 domain, linker, and CBM5 domain was visualised in UCSF Chimera (Pettersen et al. 2004). Graphics were edited using Inkscape (http://inkscape.org).

### Zoospore chemotaxis and motility

Prior to chemotaxis and motility experiments, *R. globosum* JEL800 was maintained on agar plates made with PmTG, or BBM with 5mM GlcNAc or chitin for at least 3 generations. Mature plates were sporulated as described above and 1 mL zoospores spread onto fresh media and dried to create a lawn. Lawns were incubated at 22°C for 48 hours (PmTG and GlcNAc) and 24 hours (chitin) to allow sporangia to develop and mature. For chemotaxis experiments, mature lawns were sporulated with 5mL sterile dH_2_O, and after 30 min zoospores were harvested and passed through a 10 µm sieve, enumerated using a LUNA™ Automated Cell Counter and motility checked using a haemocytometer. Only zoospore suspensions with motility >80% and ∼1x10^7^ cells/ mL were used for experiments. 2mL zoospore suspension was placed in a 50mL centrifuge tube and the top sealed with parafilm. 1 mL syringes with 21g needles containing either 150 µL 10mM GlcNAc, 10mM Glucose or 10mM H_2_O (control) were held in the zoospore suspension and incubated at 22°C in the dark. After 1 hour, the contents of the syringe was emptied into a 1.5mL tube and fixed with glutaraldehyde (final concentration of 0.5%) and zoospores enumerated using a haemocytometer. For motility experiments, zoospore lawns were sporulated with 3mL sterile dH_2_O, a 1 mL sample was taken immediately, a subsample of this was glutaraldehyde fixed to determine the starting concentration. Non-motile zoospores were enumerated using a haemocytometer every 5 min for 1 hour and % motility was calculated from the starting concentration.

## Acknowledgements

We thank Joyce E. Longcore (University of Maine) for providing *R. globosum* JEL800, now held at the Collection of Zoosporic Eufungi at the University of Michigan (CZEUM), and information on the isolation of *Podochytrium* sp. JEL0797, *Siphonaria* sp. JEL0065 and *Physocladia obscura* JEL0513. We also thank Antonia Feilden (University of York) for useful suggestions on protein modelling. NC, KB and MC were supported by the European Research Council (ERC) (MYCO-CARB project; ERC grant agreement no. 772584). DL was supported by an EnvEast Doctoral Training Partnership (DTP) PhD studentship funded from the UKRI Natural Environment Research Council (NERC grant no. NE/L002582/1).

## Notes

### Competing Interest Statement

The authors have declared no competing interest.

### Summary of Updates

We have performed additional analysis of predicted protein structure.

## References

Barr, D. J. S. 1986. Allochytridium expandens rediscovered: Morphology, physiology and zoospore ultrastructure. Mycologia 78:439–448.

Berbee, M. L., T. Y. James, and C. Strullu-Derrien. 2017. Early Diverging Fungi: Diversity and Impact at the Dawn of Terrestrial Life. Annual Review of Microbiology 71:41–60.

Ciach, M. A., J. Pawłowska, P. Górecki, and A. Muszewska. 2024. The interkingdom horizontal gene transfer in 44 early diverging fungi boosted their metabolic, adaptive, and immune capabilities. Evolution Letters.

Courtade, G., Z. Forsberg, E. B. Heggset, V. G. H. Eijsink, and F. L. Aachmann. 2018. The carbohydrate-binding module and linker of a modular lytic polysaccharide monooxygenase promote localized cellulose oxidation. The Journal of biological chemistry 293:13006–13015.

Duo-Chuan, L. 2006. Review of fungal chitinases. Mycopathologia 161:345–360.

Farrer, R. A., A. Martel, E. Verbrugghe, A. Abouelleil, R. Ducatelle, J. E. Longcore, T. Y. James, F. Pasmans, M. C. Fisher, and C. A. Cuomo. 2017. Genomic innovations linked to infection strategies across emerging pathogenic chytrid fungi. Nat Commun 8:14742.

Funkhouser, J. D., and N. N. Aronson. 2007. Chitinase family GH18: evolutionary insights from the genomic history of a diverse protein family. BMC Evolutionary Biology 7:96.

Galindo, L. J., D. S. Milner, S. L. Gomes, and T. A. Richards. 2022. A light-sensing system in the common ancestor of the fungi. Current Biology 32:3146-3153.e3143.

Gladieux, P., J. Ropars, H. Badouin, A. Branca, G. Aguileta, D. M. de Vienne, R. C. Rodríguez de la Vega, S. Branco, and T. Giraud. 2014. Fungal evolutionary genomics provides insight into the mechanisms of adaptive divergence in eukaryotes. Mol Ecol 23:753–773.

Goughenour, K. D., J. Whalin, J. C. Slot, and C. A. Rappleye. 2021. Diversification of Fungal Chitinases and Their Functional Differentiation in Histoplasma capsulatum. Mol Biol Evol 38:1339–1355.

Gow, N. A. R., J.-P. Latge, and C. A. Munro. 2017. The Fungal Cell Wall: Structure, Biosynthesis, and Function. Microbiology Spectrum 5:10.1128/microbiolspec.funk-0035-2016.

Guillard, R. R. L. 1975. Culture of phytoplankton for feeding marine invertebrates. Pages 26–60 in W. L. Smith and M. H. Chanley, editors. Culture of Marine Invertebrate Animals Plenum Press, New York.

Guy, L., J. R. Kultima, and S. G. E. Andersson. 2010. genoPlotR: comparative gene and genome visualization in R. Bioinformatics (Oxford, England) 26:2334–2335.

Jiang, Z., S. Hu, J. Ma, Y. Liu, Z. Qiao, Q. Yan, Y. Gao, and S. Yang. 2021. Crystal structure of a chitinase (RmChiA) from the thermophilic fungus Rhizomucor miehei with a real active site tunnel. Biochim Biophys Acta Proteins Proteom 1869:140709.

Joneson, S., J. E. Stajich, S. H. Shiu, and E. B. Rosenblum. 2011. Genomic transition to pathogenicity in chytrid fungi. PLoS Pathog 7:e1002338.

Jumper, J., R. Evans, A. Pritzel, T. Green, M. Figurnov, O. Ronneberger, K. Tunyasuvunakool, R. Bates, A. Žídek, A. Potapenko, A. Bridgland, C. Meyer, S. A. A. Kohl, A. J. Ballard, A. Cowie, B. Romera-Paredes, S. Nikolov, R. Jain, J. Adler, T. Back, S. Petersen, D. Reiman, E. Clancy, M. Zielinski, M. Steinegger, M. Pacholska, T. Berghammer, S. Bodenstein, D. Silver, O. Vinyals, A. W. Senior, K. Kavukcuoglu, P. Kohli, and D. Hassabis. 2021. Highly accurate protein structure prediction with AlphaFold. Nature 596:583–589.

Junges, Â., J. T. Boldo, B. K. Souza, R. L. Guedes, N. Sbaraini, L. Kmetzsch, C. E. Thompson, C. C. Staats, L. G. de Almeida, A. T. de Vasconcelos, M. H. Vainstein, and A. Schrank. 2014. Genomic analyses and transcriptional profiles of the glycoside hydrolase family 18 genes of the entomopathogenic fungus Metarhizium anisopliae. PLOS One 9:e107864.

Karlsson, M., and J. Stenlid. 2008. Comparative evolutionary histories of the fungal chitinase gene family reveal non-random size expansions and contractions due to adaptive natural selection. Evol Bioinform Online 4:47–60.

Katoh, K., and D. M. Standley. 2013. MAFFT Multiple Sequence Alignment Software Version 7: Improvements in Performance and Usability. Molecular Biology and Evolution 30:772–780.

Laundon, D., N. Chrismas, K. Bird, S. Thomas, T. Mock, and M. Cunliffe. 2022. A cellular and molecular atlas reveals the basis of chytrid development. eLIFE 11:e73933.

Laundon, D., N. Chrismas, G. Wheeler, and M. Cunliffe. 2020. Chytrid rhizoid morphogenesis resembles hyphal development in multicellular fungi and is adaptive to resource availability. Proc. R. Soc. B 287.

Laundon, D., and M. Cunliffe. 2021. A call for a better understanding of aquatic chytrid biology. Frontiers in Fungal Biology 2.

Lespinet, O., Y. I. Wolf, E. V. Koonin, and L. Aravind. 2002. The Role of Lineage-Specific Gene Family Expansion in the Evolution of Eukaryotes. Genome Research 12:1048–1059.

Liu, J., Q. Xu, Y. Wu, D. Sun, J. Zhu, C. Liu, and W. Liu. 2023. Carbohydrate-binding modules of ChiB and ChiC promote the chitinolytic system of Serratia marcescens BWL1001. Enzyme Microb Technol 162:110118.

Maillard, F., A. Kohler, E. Morin, C. Hossann, S. Miyauchi, I. Ziegler-Devin, D. Gérant, N. Angeli, A. Lipzen, K. Keymanesh, J. Johnson, K. Barry, I. V. Grigoriev, F. M. Martin, and M. Buée. 2023. Functional genomics gives new insights into the ectomycorrhizal degradation of chitin. New Phytol 238:845–858.

Medina, E. M., and N. E. Buchler. 2020. Chytrid fungi. Current Biology 30:R516–R520.

Miller, M. A., W. Pfeiffer, and T. Schwartz. 2010. Creating the CIPRES Science Gateway for inference of large phylogenetic trees. 2010 Gateway Computing Environments Workshop (GCE):1–8.

Moss, A. S., N. S. Reddy, I. M. Dortaj, and M. J. San Francisco. 2008. Chemotaxis of the amphibian pathogen Batrachochytrium dendrobatidis and its response to a variety of attractants. Mycologia 100:1–5.

Murphy, C. L., N. H. Youssef, R. A. Hanafy, M. B. Couger, J. E. Stajich, Y. Wang, K. Baker, S. S. Dagar, G. W. Griffith, I. F. Farag, T. M. Callaghan, and M. S. Elshahed. 2019. Horizontal Gene Transfer as an Indispensable Driver for Evolution of Neocallimastigomycota into a Distinct Gut-Dwelling Fungal Lineage. Appl Environ Microbiol 85.

Nagy, L. G., R. Tóth, E. Kiss, J. Slot, A. Gácser, and G. M. Kovács. 2017. Six Key Traits of Fungi: Their Evolutionary Origins and Genetic Bases. Microbiology Spectrum 5:10.1128/microbiolspec.funk-0036-2016.

Naranjo-Ortiz, M. A., and T. Gabaldón. 2019. Fungal evolution: major ecological adaptations and evolutionary transitions. Biol Rev Camb Philos Soc 94:1443–1476.

Payne, C. M., M. G. Resch, L. Chen, M. F. Crowley, M. E. Himmel, L. E. Taylor, 2nd, M. Sandgren, J. Ståhlberg, I. Stals, Z. Tan, and G. T. Beckham. 2013. Glycosylated linkers in multimodular lignocellulose-degrading enzymes dynamically bind to cellulose. Proc Natl Acad Sci U S A 110:14646–14651.

Petersen, T. N., S. Brunak, G. von Heijne, and H. Nielsen. 2011. SignalP 4.0: discriminating signal peptides from transmembrane regions. Nat Methods 8:785–786.

Pettersen, E. F., T. D. Goddard, C. C. Huang, G. S. Couch, D. M. Greenblatt, E. C. Meng, and T. E. Ferrin. 2004. UCSF Chimera--a visualization system for exploratory research and analysis. J Comput Chem 25:1605–1612.

Powell, M. J. 1983. Localization of antimonate-mediated precipitates of cations in zoospores ofChytriomyces hyalinus. Experimental Mycology 7:266–277.

Prostak, S. M., K. A. Robinson, M. A. Titus, and L. K. Fritz-Laylin. 2021. The actin networks of chytrid fungi reveal evolutionary loss of cytoskeletal complexity in the fungal kingdom. Current Biology 31:1192-1205.e1196.

Raabe, D., P. Romano, C. Sachs, H. Fabritius, A. Al-Sawalmih, S. B. Yi, G. Servos, and H. G. Hartwig. 2006. Microstructure and crystallographic texture of the chitin–protein network in the biological composite material of the exoskeleton of the lobster Homarus americanus. Materials Science and Engineering: A 421:143–153.

Richards, T. A., and N. J. Talbot. 2018. Osmotrophy. Curr Biol 28:R1179–r1180.

Roberts, C., R. Allen, K. E. Bird, and M. Cunliffe. 2020. Chytrid fungi shape bacterial communities on model particulate organic matter. Biol Lett 16:20200368.

Scholz, B., F. C. Küpper, W. Vyverman, H. G. Ólafsson, and U. Karsten. 2017. Chytridiomycosis of Marine Diatoms-The Role of Stress Physiology and Resistance in Parasite-Host Recognition and Accumulation of Defense Molecules. Mar Drugs 15.

Sipos, G., A. N. Prasanna, M. C. Walter, E. O’Connor, B. Bálint, K. Krizsán, B. Kiss, J. Hess, T. Varga, J. Slot, R. Riley, B. Bóka, D. Rigling, K. Barry, J. Lee, S. Mihaltcheva, K. LaButti, A. Lipzen, R. Waldron, N. M. Moloney, C. Sperisen, L. Kredics, C. Vágvölgyi, A. Patrignani, D. Fitzpatrick, I. Nagy, S. Doyle, J. B. Anderson, I. V. Grigoriev, U. Güldener, M. Münsterkötter, and L. G. Nagy. 2017. Genome expansion and lineage-specific genetic innovations in the forest pathogenic fungi Armillaria. Nat Ecol Evol 1:1931–1941.

Sparrow, F. K. 1960. Aquatic Phycomycetes. The University of Michigan Press, Ann Arbour.

Sprockett, D. D., H. Piontkivska, and C. B. Blackwood. 2011. Evolutionary analysis of glycosyl hydrolase family 28 (GH28) suggests lineage-specific expansions in necrotrophic fungal pathogens. Gene 479:29–36.

Stamatakis, A. 2014. RAxML version 8: a tool for phylogenetic analysis and post-analysis of large phylogenies. Bioinformatics 30:1312–1313.

Tang, B., L. Wei, W. Tang, S. Li, and R. Zhou. 2015. Effect of Linker Flexibility on the Catalytic Features of a Glycoside Hydrolase Family 45 Endoglucanase from Rhizopus stolonifer. Applied Biochemistry and Biotechnology 176:2242–2252.

Thomé, P. C., J. Wolinska, S. V. D. Wyngaert, A. Reñé, D. Ilicic, R. Agha, H.-P. Grossart, E. Garcés, M. T. Monaghan, and J. F. H. Strassert. 2023. Phylogenomics including new sequence data of phytoplankton-infecting chytrids reveals multiple independent lifestyle transitions across the phylum. bioRxiv:2023.2006.2028.546836.

Tian, W., C. Chen, X. Lei, J. Zhao, and J. Liang. 2018. CASTp 3.0: computed atlas of surface topography of proteins. Nucleic Acids Res 46:W363–w367.

Varadi, M., S. Anyango, M. Deshpande, S. Nair, C. Natassia, G. Yordanova, D. Yuan, O. Stroe, G. Wood, A. Laydon, A. Žídek, T. Green, K. Tunyasuvunakool, S. Petersen, J. Jumper, E. Clancy, R. Green, A. Vora, M. Lutfi, M. Figurnov, A. Cowie, N. Hobbs, P. Kohli, G. Kleywegt, E. Birney, D. Hassabis, and S. Velankar. 2022. AlphaFold Protein Structure Database: massively expanding the structural coverage of protein-sequence space with high-accuracy models. Nucleic Acids Res 50:D439–d444.

Venard, C. M., K. K. Vasudevan, and T. Stearns. 2020. Cilium axoneme internalization and degradation in chytrid fungi. Cytoskeleton (Hoboken) 77:365–378.

Willard, L., A. Ranjan, H. Zhang, H. Monzavi, R. F. Boyko, B. D. Sykes, and D. S. Wishart. 2003. VADAR: a web server for quantitative evaluation of protein structure quality. Nucleic Acids Res 31:3316–3319.

Yang, M., M. K. Derbyshire, R. A. Yamashita, and A. Marchler-Bauer. 2020. NCBI’s Conserved Domain Database and Tools for Protein Domain Analysis. Current Protocols in Bioinformatics 69:e90.

