## Supplementary Figures for "Evolutionary and biological mechanisms underpinning chitin degradation in aquatic fungi"

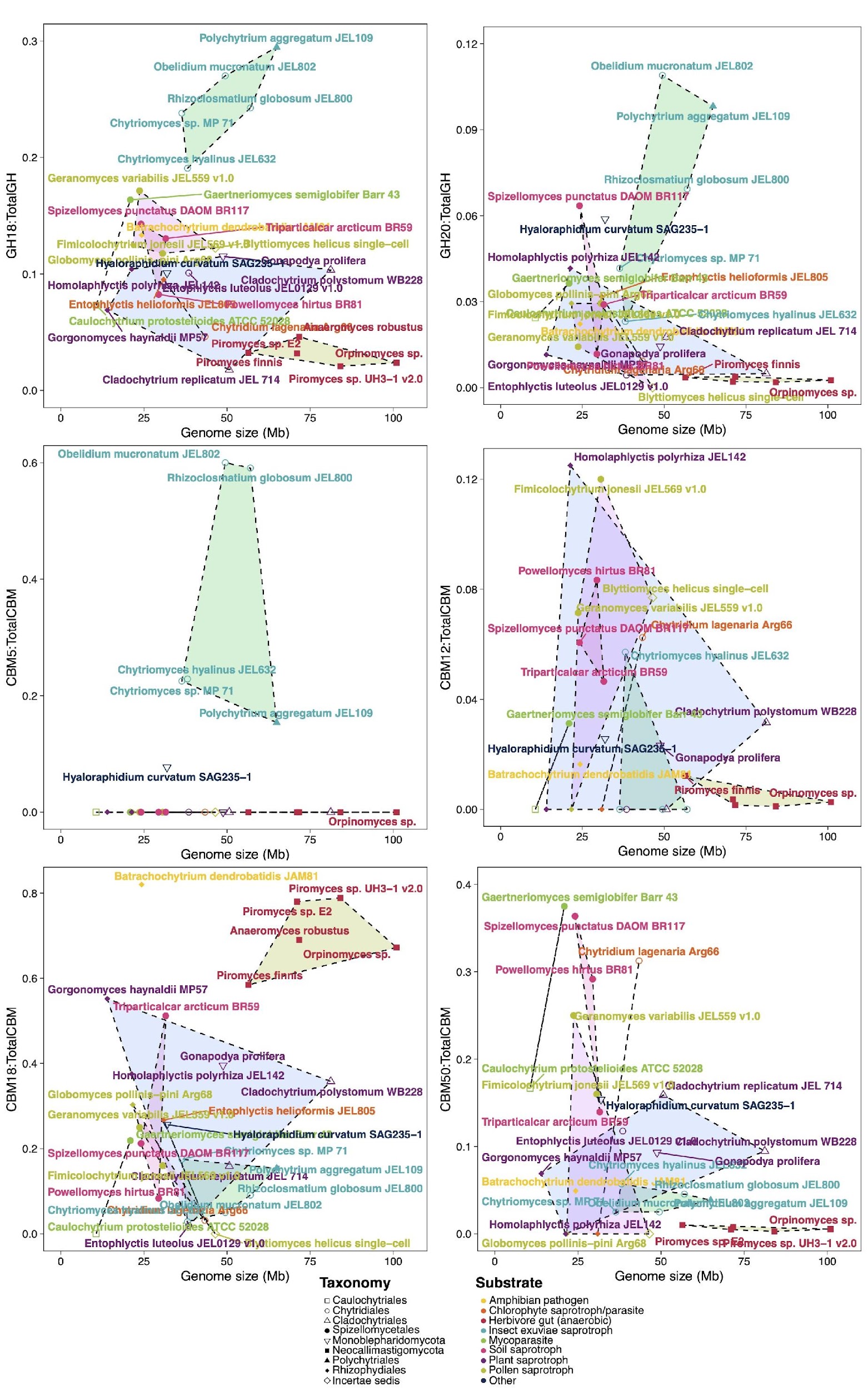

**Supplementary figure 1:** Comparison of ratios of glycoside hydrolases GH18 chitinases and GH20 β-*N*-acetyl-d-hexosaminidases with total GH encoding genes, and carbohydrate binding module CBM5, CBM12, CBM18 and CBM50 genes with total CBM encoding genes and genome sizes of 35 members of the Chytridiomycota that occupy diverse ecological niches and utilise a range of substrates.

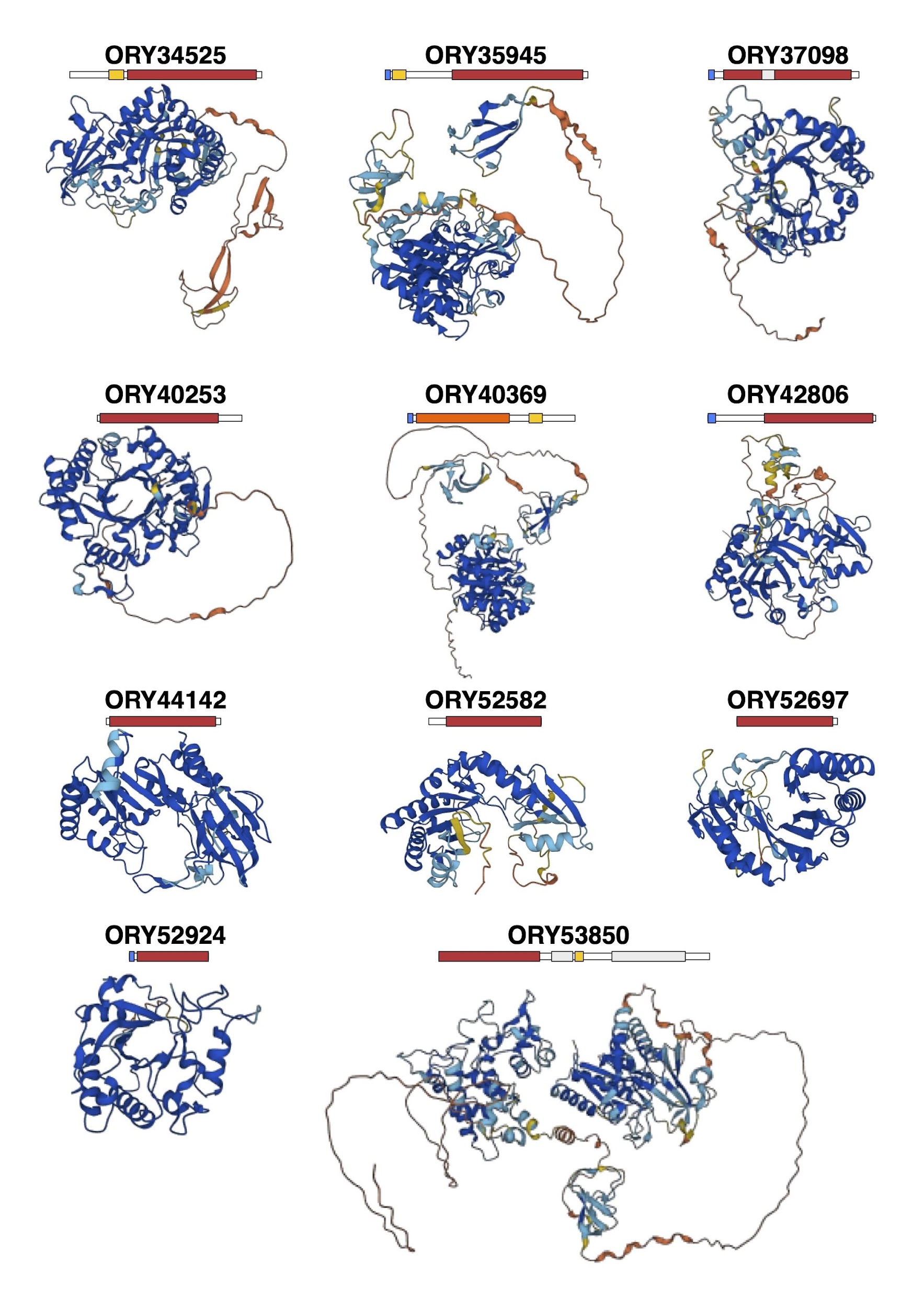

**Supplementary figure 2:** AlphaFold predicted models of secreted *Rhizoclosmatium globosum* Group AC (and other ORY40369) GH18s.

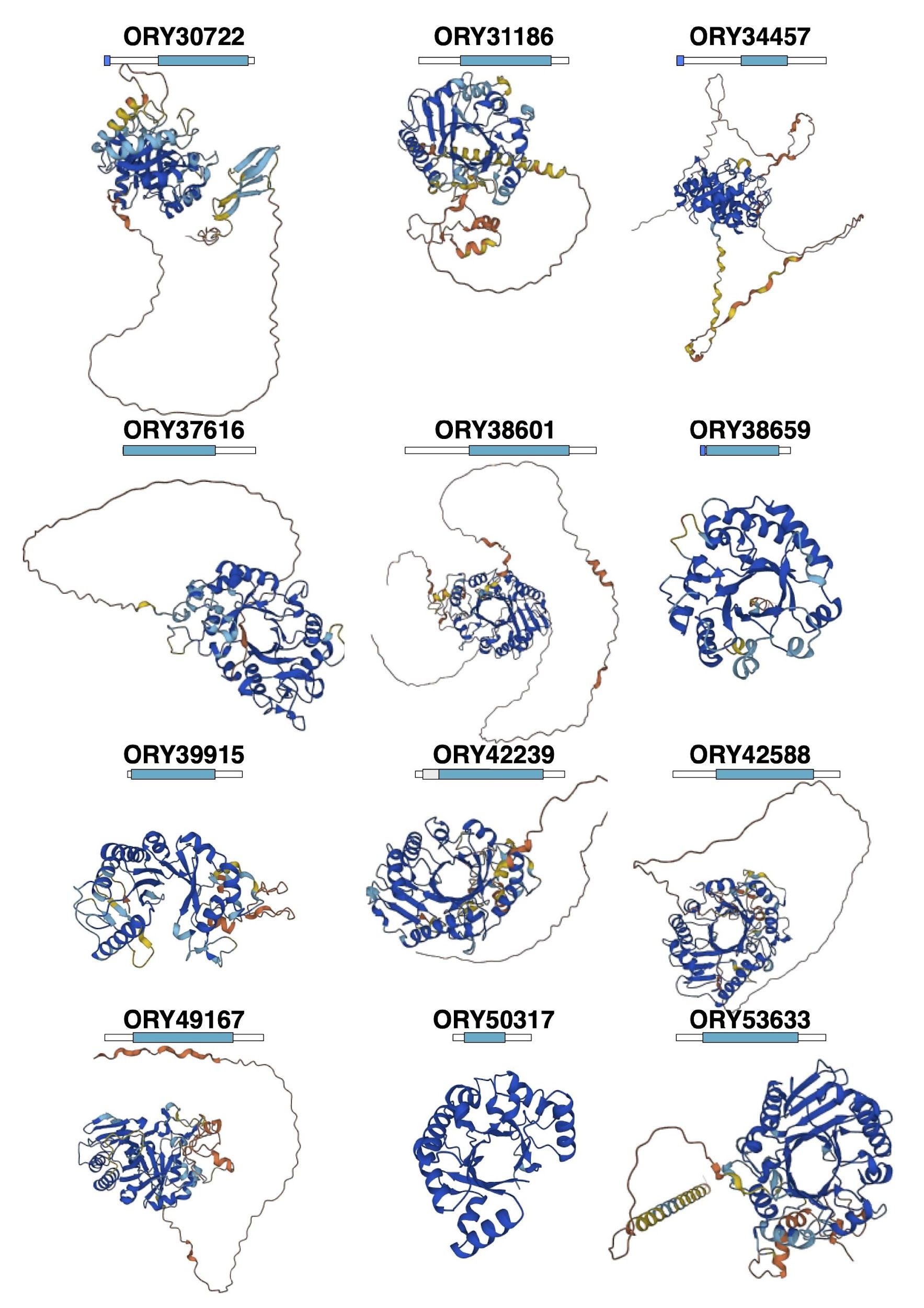

**Supplementary figure 3:** AlphaFold predicted models of secreted *Rhizoclosmatium globosum* Group B GH18s.

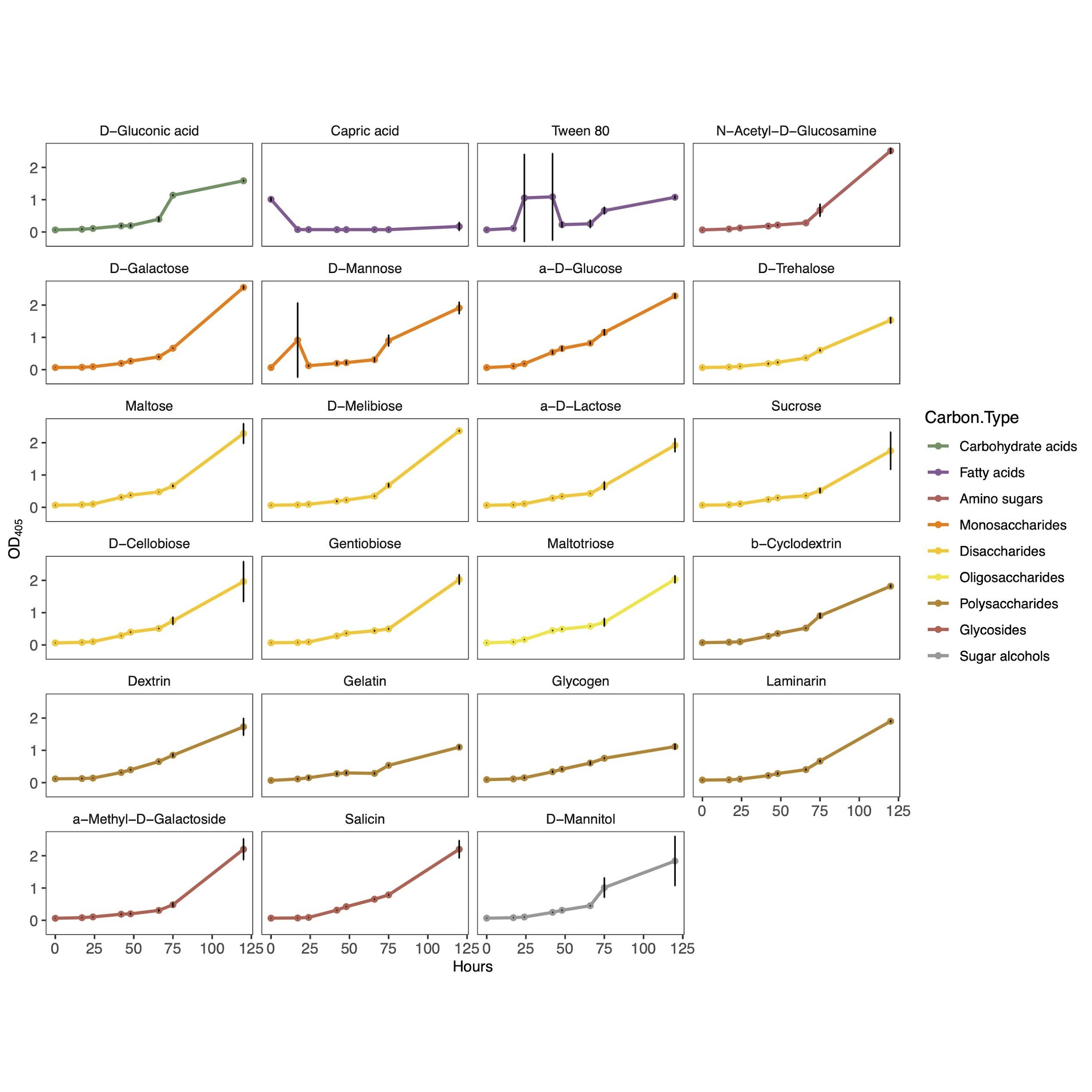

**Supplementary figure 4:** Growth curves (optical density at 405nm vs time) of significantly (p<0.05) different growth to the no C control over the whole experiment. The distribution and equality of variance of the growth data did not meet the assumptions required for ANOVA so a Kruskal–Wallis rank sum test was performed with Dunn’s post hoc analysis. Because the effect of substrates were independent to each other Dunn’s test was performed with no p value correction, then only the p values for each substrate compared to the no carbon control were extracted and the Hochberg p adjustment was applied to decrease the false discovery rate. Growth rate met the assumptions required by ANOVA so ANOVA with Dunnett’s post hoc tests were performed comparing each substrate to the no carbon control as a reference group.

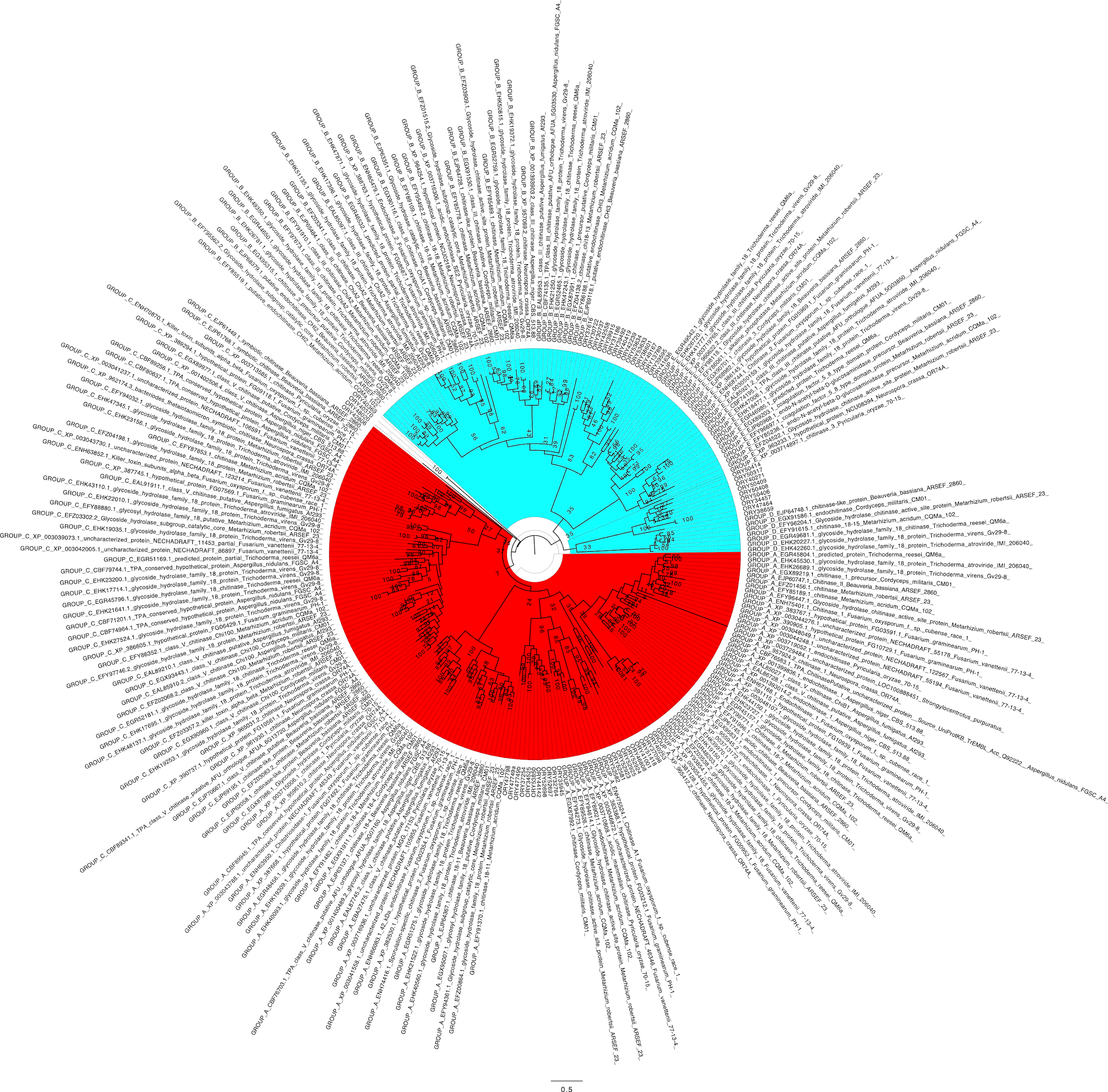

**Supplementary figure 5:** Maximum likelihood phylogenetic tree placing *Rhizoclosmatium globosum* JEL800 GH18s within the context of other fungal GH18s. Group AC are highlighted in red and Group B are highlighted in blue.
